# Synthetic biology meets proteomics: Construction of *à la carte* QconCATs for absolute protein quantification

**DOI:** 10.1101/2021.04.13.439592

**Authors:** James Johnson, Victoria M Harman, Catarina Franco, Nicola Rockliffe, Yaqi Sun, Lu-Ning Liu, Ayako Takemori, Nobuaki Takemori, Robert J. Beynon

## Abstract

We report a new approach to the assembly and construction of QconCATs, quantitative concatamers for proteomic applications that yield stoichiometric quantities of sets of stable isotope-labelled internal standards. The new approach is based on synthetic biology precepts of biobricks, making use of loop assembly to construct larger entities from individual biobricks. It offers a major gain in flexibility of QconCAT implementation and enables rapid and efficient editability that permits, for example, substitution of one peptide for another. The basic building block (a Qbrick) is a segment of DNA that encodes two or more quantification peptides for a single protein, readily held in a repository as a library resource. These Qbricks are then assembled in a one tube ligation reaction that enforces the order of assembly, to yield short QconCATs that are useable for small quantification products. However, the DNA context of the short also allows a second cycle of assembly such that five different short QconCATs can be assembled into a longer QconCAT in a second, single tube ligation. From a library of Qbricks, a bespoke QconCAT can be assembled quickly and efficiently in a form suitable for expression and labelling i*n vivo* or *in vitro*. We refer to this approach as the ALACAT strategy as it permits *à la carte* design of quantification standards.

## INTRODUCTION

Absolute quantification of proteins by mass spectrometry is typically based on the use of accurately quantified stable isotope labelled internal standards, usually peptides, as surrogates for the protein quantification. There are many ways to generate these labelled peptides, including direct chemical synthesis (AQUA peptides, [1, 2] or are derived from full length labelled proteins (PSAQ, [3–5] or shorter epitopic fragments (PrEST, [6, 7]). An additional approach is the use of QconCAT technology [4, 8, 9]. QconCATs are multiplexed protein standards for proteomics, products of artificial genes designed to encode concatamers of peptides, wherein each peptide or more commonly, a pair of peptides, is chosen to act as mass spectrometric standard(s) for absolute quantification of multiple peptides. The initial publications on QconCATs [10–12] have received over 1,000 citations and the methodology is well known and embedded in the community. At a typical size of about 60-70 kDa, a QconCAT encodes approximately 50 tryptic peptides, permitting the quantification of around 25 proteins at a ratio of two peptides per protein. The genes are then expressed as recombinant proteins in bacteria grown in the presence of SIL amino acids, usually lysine and arginine, ensuring a single labelling position for every standard tryptic peptide. Because the genes are designed *de novo*, it is feasible to introduce additional features, such as purification tags, sacrificial peptides to protect the QconCAT from exoproteolysis and peptide sequences, common to each QconCAT as a quantification standard, permitting absolute quantification of each standard within the proteomics workflow – in effect, an ‘internally standardised standard’. We have demonstrated that the QconCAT approach can be used successfully in large scale proteome quantification studies and have reported the absolute quantification of approx. 1,800 proteins in the *Saccharomyces cerevisiae* proteome [13], by far the largest absolute quantification study conducted to date. Because each peptide derived by complete excision from a QconCAT are present in equal quantities, QconCATs also have utility in determination of subunit stoichiometry of multiprotein complexes, such as the proteinaceous bacterial metabolosomes for propanediol degradation in *Salmonella* [14].

Although QconCATs have been widely adopted, their broader deployment can be challenging. First, QconCAT expression requires skills in molecular biology and facilities for bacterial expression of heterologous proteins. We have addressed this in part through the introduction of cell-free synthesis of QconCATs, which brings added advantages of concurrent, single tube synthesis that we have extended to over 100 QconCATs simultaneously, a strategy we have dubbed MEERCAT [15, 16]. Secondly, QconCATs cover a set of target proteins based on the needs of one research group, which may not always match the requirements of subsequent research groups. Thirdly, the choice of peptides is often obliged to be made without knowledge of the performance of these peptides in absolute quantification. Lastly, editing of any QconCAT, for example, the removal or addition of a single protein, has required complete resynthesis and expression of the gene.

To overcome these complications, we now introduce the concept of ‘ALACATS’ - *‘à la carte*’ QconCATs, the term reflecting the ability to design a QconCAT of any length that encodes peptides for a user-specified set of target proteins. ALACATS are assembled from ‘Qbricks’, oligonucleotides that encode (typically) two quantotypic peptides for a single target protein, together with short flanking peptides to recapitulate the correct primary sequence context, and thus normalise digestion rates. Each Qbrick (one for each target protein in the proteome) is a discrete entity, a double stranded DNA construct that can be readily synthesised, stored, catalogued and accessed to enable the synthesis of an ALACAT to order. These are the fundamental building blocks in the ALACAT workflow.

## RESULTS

### Design, synthesis and assembly of Qbricks

A Qbrick (‘quantification brick’, a type of biobrick [17]) is defined as a short, double stranded oligonucleotide that encodes two Qpeptides that are quantotypic for a single protein and is thus the smallest building block (Figure 1). The Qbrick also encodes interspersed peptide sequences that recapitulate the primary sequence context of the two peptides, thus equalising digestion rates of standard and analyte. Each Qbrick has asymmetric overhangs at each end, creating sticky ended DNA molecules that permit assembly by a strategy called ‘Loop Assembly’, driven by sequential use of two Type IIS restriction endonucleases [18]. Different unique overhang sequences (A, B, C, D, E and F) flank the Qbricks. Five Qbricks are assembled in a single reaction – the ‘odd’ cycle [18]. The annealing during assembly maintains the reading frame through the QconCAT, adding two amino acids to the interspersed linker with little or no effect on the peptides generated from the QconCAT (Figure 1a). These *short QconCATs*, assemblies of five Qbricks that encode 10 peptides, are perfectly useable when expressed as a five-target protein standard suitable for small, focused studies. A short QconCAT, containing 10 Qpeptides (from five Qbricks), interspersed linker peptides, quantification and purification peptides as well as suitable sacrificial sequences at either end, totals approximately 170-220 amino acids, of a size suitable for expression and deployment.

**Figure 1.**
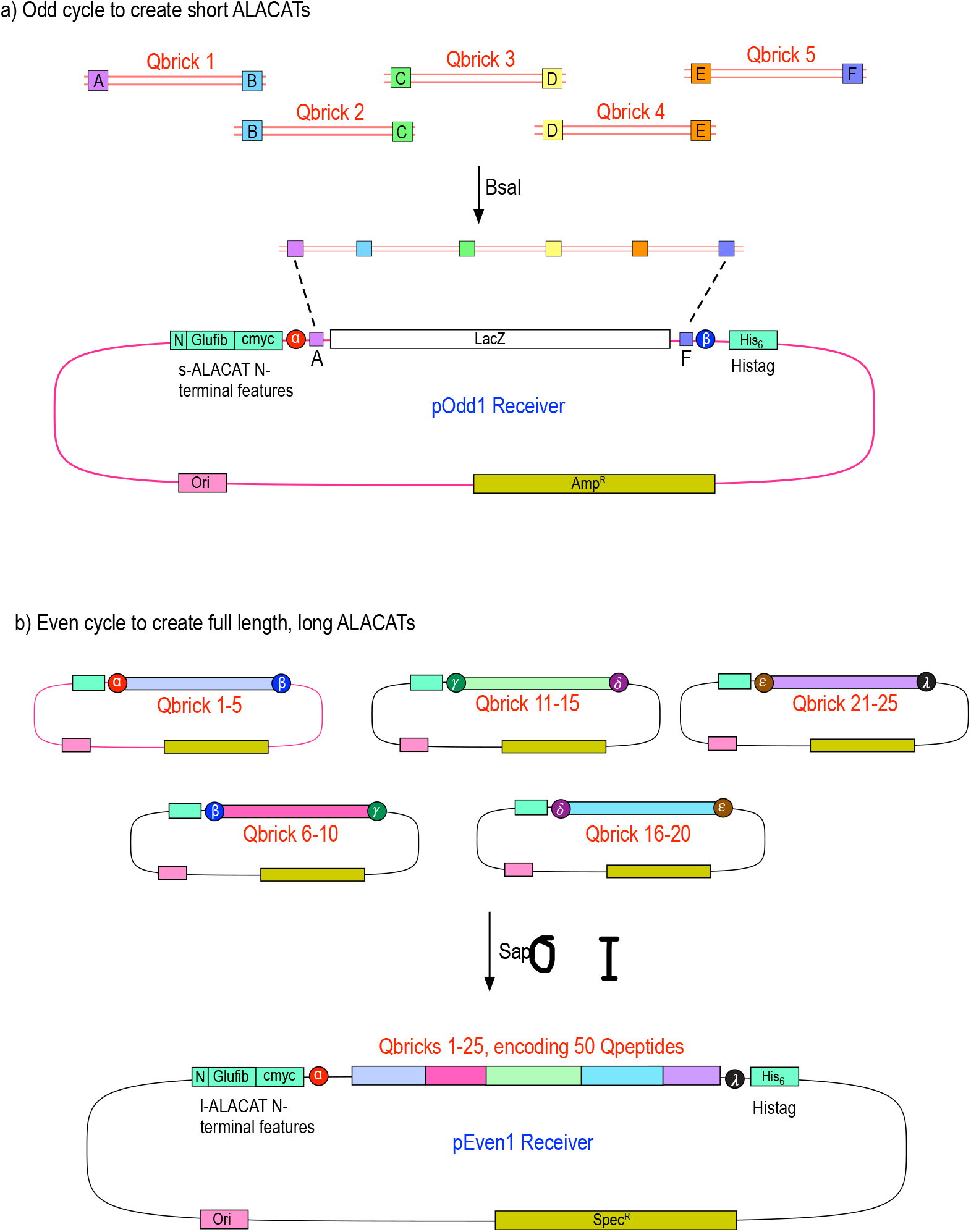
Overall strategy for building block assembly of short and long ALACATs. The smallest unit of an ALACAT is a single double stranded oligonucleotide encoding peptides (one, two, more) for quantification of a single target protein, flanked on either side by tripeptides that preserve the natural primary sequence context. These oligonucleotides include linker regions compatible with a Type IIS restriction enzyme (*Bsa*I) that allows all five Qbricks to be assembled in the correct order in a single ligation reaction (panel a, odd cycle assembly) to form short ALACATs. In turn, these short ALACATs contain DNA sequences that are compatible with a second Type IIS restriction enzyme (*Sap*I) and can be similarly assembled in a one tube reaction to create a long ALACAT (panel b, even cycle assembly). The vectors for the odd and even cycles both include in frame fusions to glu-fibrinopeptide and c-myc encoding regions (quantification) and a hexahistidine tag (purification).

For more wide-reaching quantification studies, individual short ALACATs are subsequently concatenated in a second reaction. The initial 5-Qbrick constructs are cloned into plasmids that introduce a second set of six overhanging linker sequences, distinct from those used in the ‘odd’ cycle. These linkers (‘even cycle’, α, β, γ, δ, ε, λ; Figure 1b) allow assembly of the five short ALACATs into a complete, ‘*long ALACAT*’, capable of encoding Qpeptides for approximately 25 proteins. These QconCATs would be 75-90 kDa (relatively shorter because of the single instances of N-terminal and C-terminal features), typical for cell-free or bacterial expression. Of course, any variant, from two to five short ALACATs, encoding quantification standards for any number of proteins between 5 and approximately 30 is possible using this approach. This greatly expands the flexibility of the QconCAT approach. The sequences of the constructs used in this paper, and the cloning syntaxes, are provided in Supplementary Sequence File A.

As proof of concept, we built an ALACAT from a series of Qbricks encoding standards for 25 human plasma proteins (Supplementary Table 1). We first assembled five short ALACATs and then, in turn, assembled these into a long ALACAT. Each short ALACAT was expressed independently in a wheat germ cell free system (CFS), as well as the long ALACAT and yields of all were high (Figure 2a). Typically, yields were of the order of 500 pmol, which is a substantial quantity for LC-MS/MS based quantification (typically, a single LC-MSMS run would require 50 fmol on column). The expressed short and long QconCATs were then digested with trypsin and analysed by LC-MS/MS (Figure 2b). Further information on analysis of the short and long ALACATs is provided in Supplementary Material.

**Figure 2.**
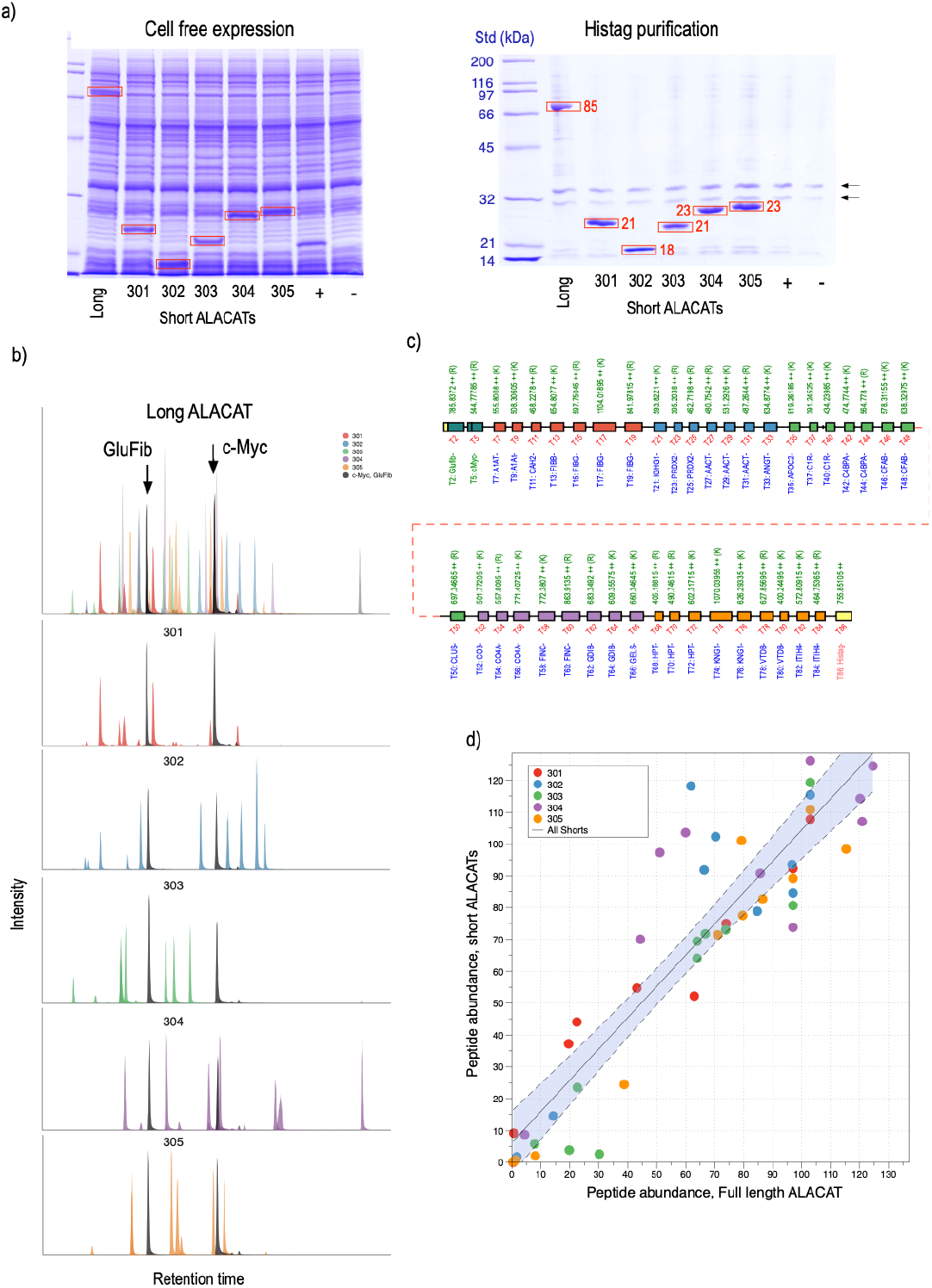
Construction of a human plasma protein QconCAT using the ALACAT strategy. A series of Qbricks were designed, each encoding two peptides for each of 25 plasma proteins. Groups of five were assembled in an odd cycle reaction to short ALACATs (301-305) that were expressed *in vitro* using wheat germ lysate and purified by virtue of their hexahistidine tag. In addition, the short ALACATs were assembled in an even cycle reaction into a single long ALACAT that was also expressed and purified (panel a). Each of the ALACATs were then digested and analysed by LC-MSMS; all peptides were detected (panel c, infilled peptides are those visible by LC-MS/MS (panel b); different colours define the short ALACAT origins of the peptides in the long ALACAT). Using common peptides (glu fibrinopeptide and c-myc epitope) as normalisation controls, the peak areas for peptides in short ALACATs were compared to the areas of the same peptides in the long ALACATs (panel d)

Each QconCAT contained two common peptides - the glu fibrinopeptide (EGVNDNEEGFFSAR) that we have used previously for quantification of the QconCAT [8, 13] and a second peptide derived from the common c-myc peptide (LISEEDLGGR) to give a tag for monitoring expression by western blotting if necessary. We were able to use these two peptides to assess the consistency of the intensity of the quantification peptide, whether in long or short ALACATs (Figure 2c). The correlation was extremely high, confirming that the peptides were cleaved and released similarly, irrespective of the nature of the ALACAT.

We also demonstrated the synthesis of ALACAT in a *E. coli* cell-free system. The *E. coli* system couples transcription and translation in a single tube, which allows us to skip the *in vitro* transcription reaction required in the wheat cell-free system. In this study, we set up a small-scale reaction system using a microdialysis device (Figure 3). All prepared ALACAT genes were successfully synthesized in this system), and the efficiency of ^13^C/^15^N incorporation into their lysine and arginine residues was more than 99 %.

**Figure 3.**
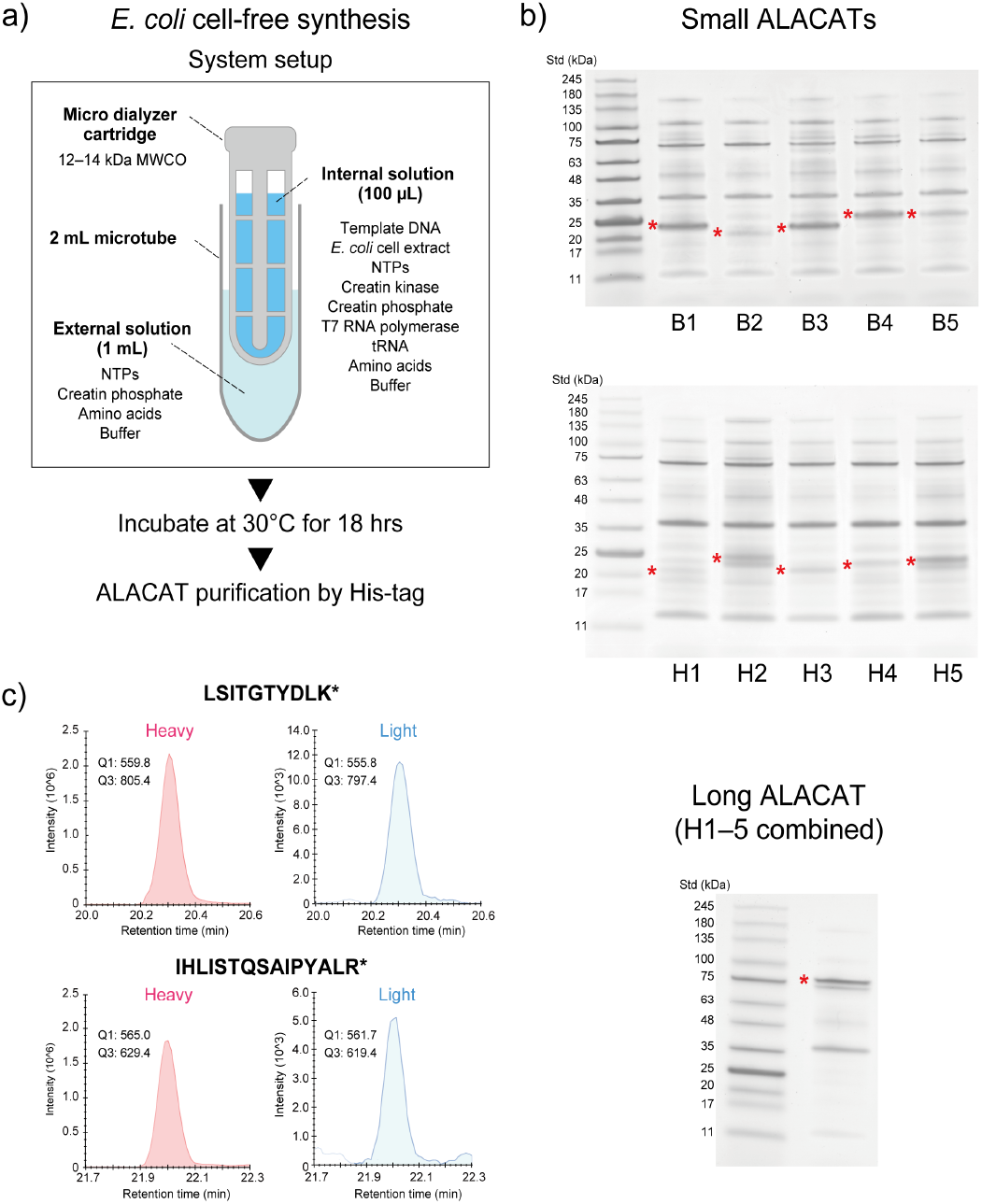
ALACAT synthesis by *E. coli* cell-free system. Experimental design for cell-free synthesis (panel a). Ten small ALACATs (belonging to two series; ‘B’ and ‘H’) and one long ALACAT were synthesized in a small-scale cell-free synthesis system using a microdialysis device (panel b). Lysine and arginine residues of the synthesized ALACATs were labelled with 13C/15N. Panel b) Representative SDS-PAGE separation images of the synthesized ALACATs. His-tag purified ALACATs were separated by NuPAGE 4–12 % gradient gels and visualized by CBB staining. Asterisk: ALACAT band. Panel c) Selective reaction monitoring (SRM) analysis of ALACAT peptides. The efficiency of stable isotope incorporation was estimated by SRM for two tryptic digested peptides derived from one ALACAT.

### Editability of ALACATs

One of the advantages of the ALACAT approach is the introduction of straightforward editability of the construct. Previously, there was no simple route to exchange one peptide for another without extensive resynthesis of the gene. However, with ALACATs, the editability simplifies the introduction of changes in the sequence and embedded peptides. This editing process can take place at two levels. First, individual peptides can be replaced in Qbricks, and a new short ALACAT could be constructed. The only new DNA required would be the sequence of the Qbrick. Alternatively, an entire short ALACAT could be exchanged, replacing multiple peptides in a single process. This might be of particular value, for example, in a multi-species construct, if some short ALACATs contained species-specific peptide sequences, and others contained sequences that were identical in both species. A simple switch from species A to species B would only require exchange of the relevant short ALACAT in the one-step, even cycle ligation reaction. Further, about 10% of all traditionally synthesised QconCATs failed to express in bacteria [16] and the ability to quickly create a large set of rearrangements of different Qbricks or short ALACATs would be able to deliver a library of ALACATs, with equivalent function, that could be quickly screened for expression potential. Alternatively, this type of combinatorial synthesis could be used to explore adjacency and proximity effects. To test this possibility *in extremis* we therefore initiated a ‘one pot’ combinatorial ligation of two families of Qbricks, or two families of short ALACATs.

We tested both levels of editability using the two ALACAT series (B, plasma and H for analysis of the stoichiometry of a metabolic compartment; Supplementary Sequence File A) described above. First, we demonstrated the ease of exchange of short ALACATs by building a combinatorial series of long ALACATs created from random introduction of appropriate short ALACATs – the ‘even’ cycle. Each position in the long ALACAT could contain a short ALACAT from either the B or H series. Rather than create one editing reaction to prove the swap of one short ALACAT for another, we took a different approach and set up a single reaction, in which we mixed ten short ALACATs derived from the two different families, prefixed B and H, such that B1 and H1 would share common *Sap*I overhang sequences and similarly, the other four pairs (H2/B2 to H5/B5). Thus, short ALACATs H1 and B1 would represent a binary choice at position one. In this assembly, a random ligation process would generate 2^5^ = 32 different combinations, from H1:H2:H3:H4:H5 through, for example, H1:H2:B3:B4:B5 etc. to B1:B2:B3:B4:B5. After the single tube ligation, 81 colonies were picked and the ligation product DNA was sequenced. Of the 32 combinations that could have been synthesised, we detected 26 different short ALACATs (80% of all possible different combinations, Figure 4, Supplementary Sequence File B), confirming the ease of editing and reconstruction of new short ALACATs. There was no indication of any systematic bias in the selection of one or the other sets of Qbricks, establishing the ease of random ligation.

**Figure 4.**
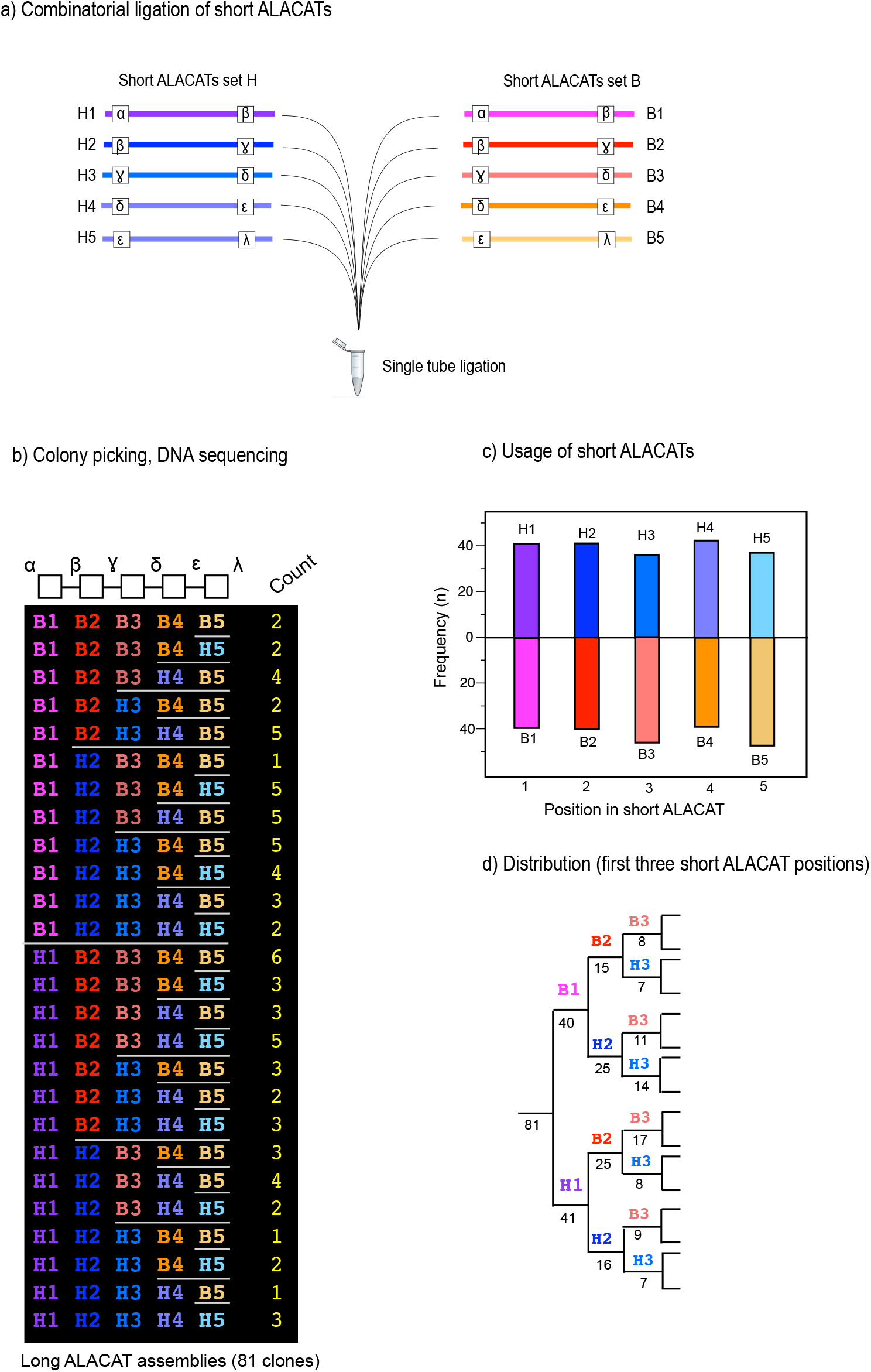
Combinatorial assembly of short ALACATs into long ALACATs. To test the ease of editability and swapping of short ALACATs into long ALACATs, we created a ligation reaction in which there were two choices of short ALACAT at each of four or five positions (panel a). The reaction products were cloned and a total of 81 clones were selected for DNA sequencing, to establish the composition of the long ALACATs (from either the B or H series) were incorporated (panel b). There was an even representation of the short ALACATs across the entire structure (panel c) and the evenness of the creation of the products is evidenced by the split of the products through the first three positions (panel d).

To assess the equivalent combinatorial substitution of Qbricks, we performed essentially the same experiment in an ‘even’ cycle but with two set of Qbricks from the two families (B and H), again picking multiple clones from a single tube ligation reaction. To increase complexity, we created further potential by providing H4 and B5 with two assembly contexts (Figure 5, Supplementary Sequence File C). After assembly, multiple ALACAT clones were picked and sequenced. From this experiment, 26 unique short ALACATs were constructed, spread across 65 sequences that were sampled. Of these 65, seven were long variants of five Qbricks (made possible by our construction strategy) but the majority comprised assemblies of four Qbricks, a total of 19 combinations from a set of 24 possibilities were recovered. Further, 16 were unique, two were replicated once, three occurred thrice, two were four-fold, up to one assembly that was sequenced in 16 (approx. 25%) of the clones. It is possible that this bias reflected differences in the relative concentrations of the input DNA sequences, which would allow for the possibility of a degree of tuning of the system to preferred assemblies.

**Figure 5.**
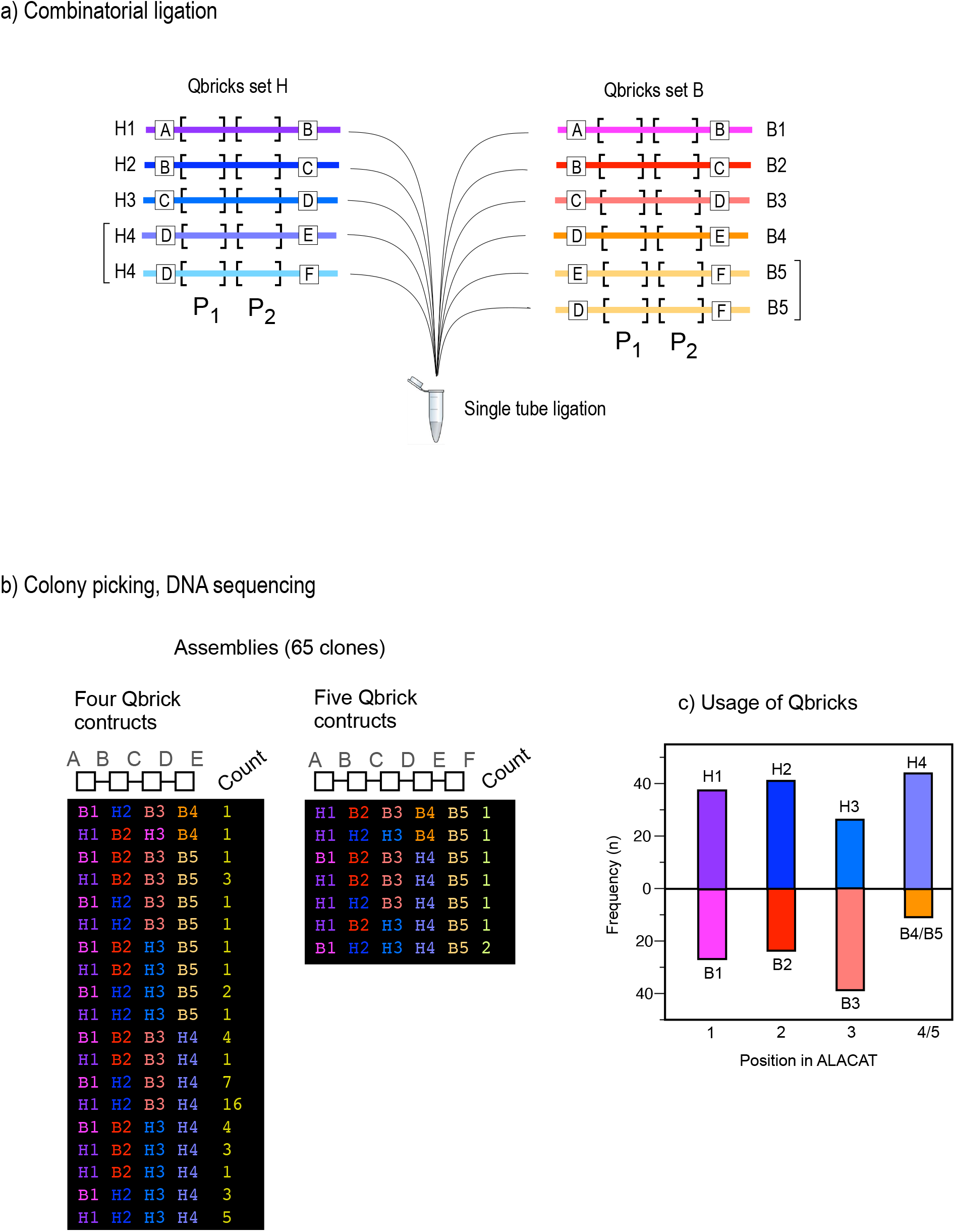
Combinatorial assembly of Qbricks into short ALACATs. To test the ease of editability and swapping of Qbricks into short ALACATs, we created a ligation reaction in which there were two choices of Qbrick at each of four or five positions (panel a), encoding pairs of quantotypic peptides P1 and P2. The reaction products were cloned and a total of 65 clones were selected for DNA sequencing, to establish the composition of the long ALACATs (from either the B or H series) were incorporated (panel b). There was a reasonably even representation of the Qbricks across the entire structure (panel c).

## Discussion

The Qbrick concept has multiple advantages over traditional QconCAT gene synthesis and expression. First, the individual Qbrick DNA oligonucleotides can be drawn from an ever-expanding library, stored at the point of synthesis, and thus, the clustering of Qpeptides into QconCATs becomes a late decision, driven by the interests of specific users and programmes. Secondly, ALACATs can be designed and delivered at any size, according to the focus and depth of individual quantitative proteomics studies. For example, two or three short ALACATs can be combined to form highly efficient QconCATs of intermediate size. Lastly, if specific peptides are suboptimal for a particular mass spectrometric approach, often an unknown factor before the construct is made, it is trivial to replace one Qbrick, build a new short ALACAT, and if required, subsequently assemble the new short ALACAT into the full length ALACAT, both steps being single-tube reactions. This is an exciting improvement in QconCAT deployment, providing an efficiency and flexibility which, coupled with cell-free synthesis and MEERCAT, means that large scale absolute proteome quantifications are now eminently feasible, sustainable and modifiable. Moreover, many stages of the ALACAT workflow are suitable for delivery through laboratory automation, reducing the need for human intervention. The Qbrick approach means that it would be possible to create an ever-expanding resource of Qbrick DNA (in the form of double stranded oligonucleotides) that could be assembled ‘to order’ in response to requests by any research group. There would be no reliance on prior clustering of peptides, and the assembly would be a trivial additional step. Moreover, the ability to ‘swap out’ specific Qbricks without having to redesign and build the QconCAT from scratch means that problematic peptides will be rapidly expunged from the resource (Figure 6). The advantages of having ‘editable QconCATs’ cannot be overstated. This added flexibility in standard design and optimisation, coupled with ever increasing selectivity and sensitivity of LC-MS/MS platforms, makes absolute quantification of part, or even all, of a proteome increasingly feasible.

**Figure 6.**
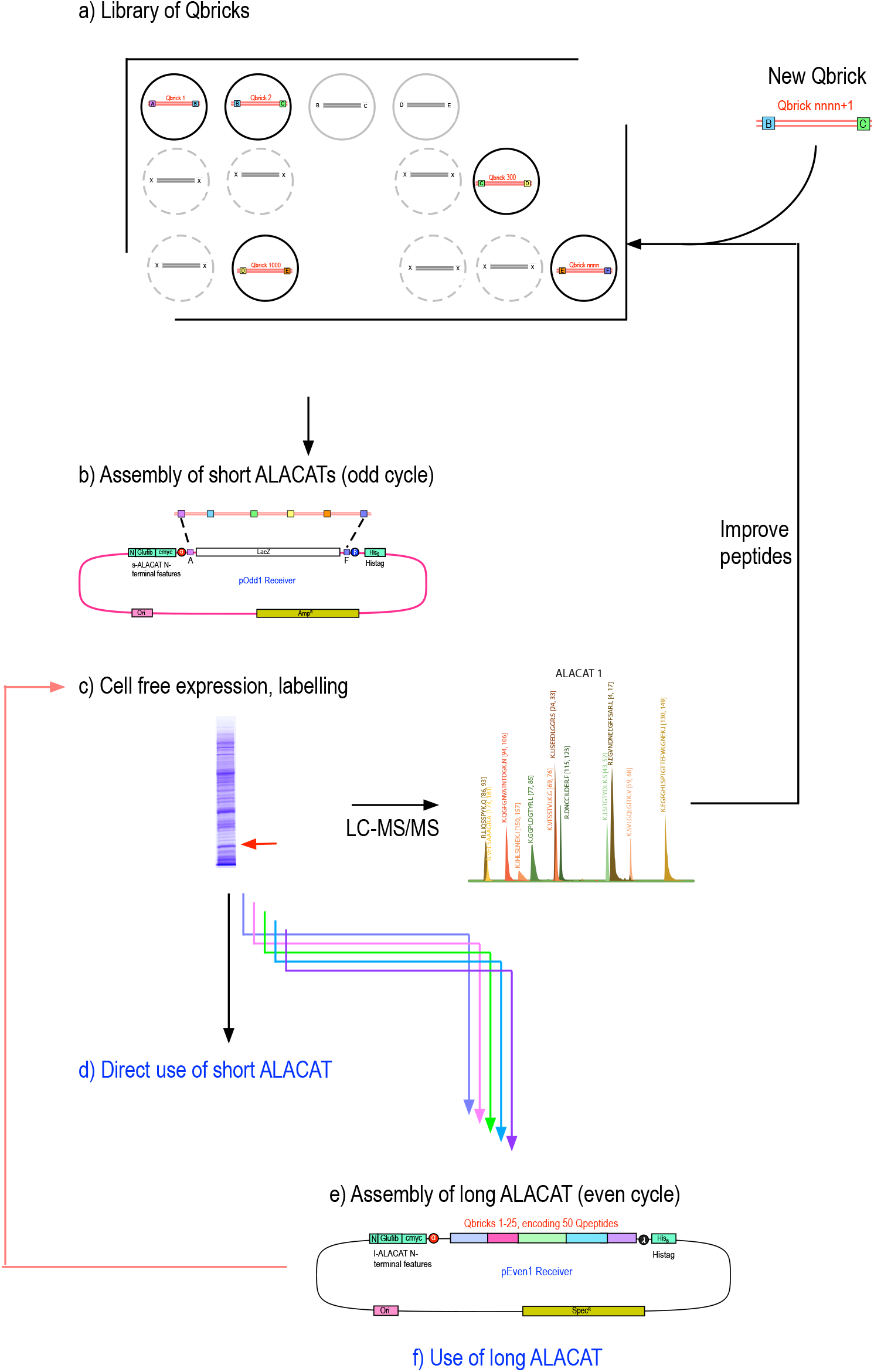
A model for the creation of an ALACAT resource. From a library of Qbricks (that may include redundant sequences to increase the choice of quantotypic peptides for specific proteins) assembly into short ALACATs would provide a test system to assess the suitability of peptides for quantification. Once the short ALACATs are optimised, sets of them could subsequently be assembled into long ALACATs. Long ALACAT DNA can be readily used to drive protein synthesis in the end user laboratory, labelled as appropriate.

Lastly, QconCATs, assemblies of peptides generated by proteolysis, are a simple route to the generation of stoichiometric quantities of a set of peptides that can be used for purposes other than absolute quantification, such as instrument quality control or calibration of retention time index ([19–23]).The combinatorial experiments described in this paper, for example, create the ability to build a large number of different combinations of peptides from a common library, and could be used in the understanding of local influences on ionisation, or even to test the emergent methods for prediction of precursor or product ion intensity [24–29]. The more straightforward the production methodology, the more likely tests of such predictive methods can be created.

## Supporting information

Supplementary Sequence File A

Supplementary Sequence File B

Supplementary Sequence File C

## Acknowledgments

This work was supported by the Biotechnology and Biological Sciences Research Council (BB/S020241/1, RJB). This work was supported by the Biotechnology and Biological Sciences Research Council (BB/R003890/1, BB/M024202/1, BB/R01390X/1, L-NL) and the Royal Society (URF\R\180030, RGF\EA\181061, RGF\EA\180233, L-NL). We are grateful to Dr Philip Brownridge for excellent instrumentation support.

## EXPERIMENTAL

### Materials and reagents

All enzymes, competent cells and manual DNA purification kits were purchased from New England Biolabs (Hitchin, UK), all oligonucleotides were purchased desalted and lyophilised from Integrated DNA Technologies BVBA (Leuven, Belgium) or Eurofins Genomics (Ebersberg, Germany). All bacterial media and antibiotics were purchased from Formedium Ltd (Hunstanton, UK).

### Production of pOdd and pEven acceptor vectors

Plasmid pEU01-MCS (CellFree Sciences, Ehime, Japan) was domesticated via site-directed mutagenesis to remove unwanted *Bsa*I and *Sap*I restriction sites. pOdd vectors were produced by inserting a lacZ cassette with *Sap*I and *Bsa*I sites as indicated with appropriate syntaxes flanked N-terminally by GluFib and Myc tag linkers and C-terminally with 6x His-Tag and stop codons. pEven vectors were produced similarly but the Amp^R^ gene was exchanged for Spec^R^ gene amplified from pGM134_1 and cloned via an NEBuilder (NEB, UK) reaction. All lacZ cassettes were synthesised by Twist Bioscience (San Francisco, USA) and cloned as single fragments into modified pEU01-MCS via NEBuilder, producing pOdd vectors pGM247_2 – 6 and pEven vectors pGM247_8 – 12.

### Design of oligonucleotides and production of QBrick DNA Blocks

QBrick peptide sequences were reverse translated using Geneious software (Biomatters Ltd), set up to use the *Escherichia coli* K12 codon usage table [30] and to avoid internal restriction sites of *Bsa*I, *Sap*I, *Bbs*I and *Bsm*BI and >5 nt homopolymers. These were converted to overlapping oligonucleotides (T_ann_ approx. 60°C). Required 5’ overhangs for *Bsa*I or *Sap*I recognition sequences and molecular syntaxes were then added. Pairs of overlapping oligonucleotides were mixed at 2.5 µM ea. (final conc.) in Q5 2x mastermix in 20 µl total reaction. These were annealed and extended using the following thermocycler parameters: 98°C for 60 s followed by five cycles of 98°C for 10 s, 60°C for 30 s, 72°C for 15 s and a final incubation at 72°C for 60 s. These reactions were diluted 1:100 in water before added to cloning reactions below (approx. 25 fmol/µl).

### Odd level cloning reactions (Short ALACATs)

Required QBrick blocks (∼25 fmol·µL^-1^) were reacted with empty pOdd vector in 10 µL total as follows: 0.5 µL each QBrick block, 10 ng pOdd vector, 1 µL T4 DNA Ligase Buffer, 0.5 µL T4 DNA Ligase (400 U·µL^-1^), 0.5 µL *Bsa*I (10 U·µL^-1^). These were incubated with the following conditions: 37°C for 10 min, followed by 50 cycles of 37°C for 1 min and 16°C for 1.5 min before final 50°C for 5 min. 1 µL of the reaction was transformed into 25 µL of NEB 5-alpha chemically competent cells. 20% were plated onto LB agar plates containing 100 µg·mL^-1^ Carbenicillin, 20 µg·mL^-1^ X-Gal and 100 µM IPTG for blue/white selection. White colonies were then screened by colony PCR using universal screening primers (Forward: 5’-TAACCACCTATCTACATCACC-3’ and Reverse: 5’-CGAGCTCGAGAACTAGTGAT-3’). PCR products were analysed using QIAxcel DNA Screening Gel automated electrophoresis (QIAGEN, Manchester, UK). Correct PCR products were cleaned with ExoCIP and Sanger sequenced (Source Bioscience Ltd, Nottingham, UK) to confirm the sequence.

### Even level cloning reactions (Long ALACATs)

Required pOdd clones (10 ng·µL^-1^) were reacted with empty pEven vector in 10 µL total as follows: 0.5 µL each pOdd clone, 10 ng pEven vector, 1 µL 10x T4 Ligase Buffer, 0.5 µL T4 DNA Ligase (400 U·µL^-1^), 0.5 µL *Sap*I (10 U·µL^-1^). These were incubated at 37°C for 120 min. 1 µL of the reaction was transformed into 25 µL of NEB 5-alpha chemically competent cells. 20% were plated onto LB agar plates containing 50 µg·mL^-1^ Spectinomycin, 20 µg·mL^-1^ X-Gal and 100 µM IPTG. White colonies were then screened by colony PCR using universal screening primers (Forward: 5’-TAACCACCTATCTACATCACC-3’ and Reverse: 5’-CGAGCTCGAGAACTAGTGAT-3’). PCR products were analysed using QIAxcel DNA Screening Gel electrophoresis.

### Cell-free expression of short and long ALACATs

For each ALACAT, 2 μg DNA in pEU-E01 vector (CellFree Sciences Co., Ltd, Japan) was used for a single expression reaction. Synthesis was completed in 240 μL scale using WEPR8240H full Expression kit (2BScientific Ltd, UK). A positive control (pEU-E01-DHFR coding dihydrofolate reductase gene derived from *E. coli*) and negative control (pEU-E01-MCS empty vector) were used, both supplied with the kit. Full kit instructions were followed, including preparation of WEPRO8240H aliquots and 2 x SUB AMIX reagent. The Transcription Mix for each expression was prepared with 20 U RNase inhibitor, 20 U SP6 RNA Polymerase, 50 nmol NTP mix and a 0.2 x dilution Transcription Buffer. DNA for the ALACAT or controls, and nuclease-free water, were added to a final volume of 20 μL. The transcription reaction occurred over six hours at 37 °C and the resulting mRNA was stored briefly at room temperature before transcription.

A 1 x SUB AMIX was prepared with a 0.5 x dilution of 2 x SUB AMIX into nuclease-free water and 60 nmol of each of the standard 20 amino acids (R, K, A, N, D, C, E, Q, G, H, I, L, M, F, P, S, T, W, Y, V), with substituted stable isotope labelled [^13^C_6_],[^15^N_4_] arginine and [^13^C_6_],[^15^N_2_] lysine (CK Isotopes Ltd, UK). The Translation Mix for each expression was prepared with 12 nmol of each of the same standard 20 amino acids, including ^13^C_6_ ^15^N Arg and Lys, combined with 0.8 μg creatinine kinase, 10 μL WEPRO8240H wheat germ lysate, 0.5 x dilution of 2 x SUB AMIX, and 10 μL of mRNA for each ALACAT or standard. A 96-well plate was prepared with 200 μL of 1 x SUB AMIX in each well. The Translation Mixture was carefully pipetted beneath the SUB AMIX in each well to form a bilayer. The plate was sealed and incubated at 16 °C for 16 hours.

*E. coli* cell-free synthesis was performed using a Musaibou-Kun protein synthesis kit (Catalog #A183-0242, Taiyo Nippon Sanso Corporation, Tokyo, Japan). For ALACAT synthesis, an amino acid cocktail with lysine and arginine universally labeled with ^13^C and ^15^N (Catalog # A91-0128, Taiyo Nippon Sanso Corporation) was used. All synthetic reactions were performed using an Xpress micro-dialyzer MD100 with molecular weight cut-off of 12–14 kDa (Scienova, Spitzweidenweg, Germany) inserted into a 2 mL microtube. Before synthesis, 825 μL of the outer solution was mixed with 75 μL amino acid cocktail and 100 μL distilled water, incubated at 30 °C, and added to the outside of the dialysis unit at the start of synthesis. Then, 77.5 μL of the internal solution for synthesis was mixed with 10 μL template DNA (50 ng/μL), 7.5 μL amino acid cocktail, and 5 μL distilled water, and added to the dialysis unit. The synthesis reaction was carried out at 30 °C for 18 hours. After the synthesis was completed, all the solution in the dialysis unit was collected into a new tube.

### ALACAT purification

Note that the positive control used for expression does not have a hexa-histidine tag and therefore both controls were used as negative controls in this next stage of the protocol. The 240 μL contents of each individual well of the 96 well plate was transferred to a low binding tube (Biotix Inc., US). This was then combined with 400 μL Bind Buffer pH 7.4 (20 mM sodium phosphate, 0.5 M sodium chloride, 20 mM imidazole, 6 M guanidine hydrochloride), and incubated at room temperature for one hour using a rotor mixer, before addition of 10 μL Ni Sepharose suspension (GE Healthcare Ltd, UK) and a further one hour incubation. Centrifuge filters (Corning Costar Spin-X 0.45 um pore size cellulose acetate membrane, Merck, UK) were washed once with 750 μL Bind Buffer and centrifuged, before addition of the sample and Ni Sepharose, and further centrifugation; all centrifuge steps used 6,000 x *g* 2 minutes 4 °C. This was followed by three further washes by centrifugation with Bind Buffer; two 400 μL washes and one 200 μL wash. Sample was eluted by centrifugation from the resin with two additions of 15 μL Elution Buffer pH 7.4 (20 mM sodium phosphate, 0.5 M sodium chloride, 1 M imidazole, 6 M guanidine hydrochloride), after each addition the resin and buffer were agitated to mix before centrifugation.

The final 30 μL elution was transferred to a low binding tube for protein precipitation. To each tube 600 μL HPLC grade methanol (Fisher Scientific Ltd, UK) was added and mixed well before addition of 150 μL chloroform and 400 μL HPLC grade water (VWR International, UK) to precipitate proteins. Following centrifugation at 13,000 x *g* for 3 minutes a bilayer was formed, the uppermost layer of which was carefully removed. A further 600 μL methanol was added and gently mixed by inversion. After a second centrifugation step the majority of the liquid was removed and discarded, with the remaining liquid allowed to evaporate. The precipitate was resuspended in 30 μL 25 mM ammonium bicarbonate, with 0.1 % (w/v) RapiGest™ SF surfactant (Waters, UK) and protease inhibitors (Roche cOmplete™, Mini, EDTA-free Protease Inhibitor Cocktail, Merck, UK). Before tryptic digestions the protein concentration of each sample was determined using a NanoDrop Spectrophotometer (ThermoFisher Scientific, UK).

### Tryptic digestion

For digestion, 0.5 μg protein for each was treated with 0.05 % (w/v) RapiGest™ SF surfactant at 80 °C for 10 minutes, reduced with 4 mM dithiothreitol (Melford Laboratories Ltd., UK) at 60 °C for 10 minutes and subsequently alkylated with 14 mM iodoacetamide at room temperature for 30 minutes. Proteins were digested with 0.01 μg Trypsin Gold, Mass Spectrometry Grade (Promega, US) at 37 °C overnight. Digests were acidified by addition of trifluoroacetic acid (Greyhound Chromatography and Allied Chemicals, UK) to a final concentration of 0.5 % (v/v) and incubated at 37 °C for 45 minutes before centrifugation at 13,000 x *g* 4°C to remove insoluble non-peptidic material.

### LC-MS/MS

Samples were analysed using an UltiMate™ 3000 RSLCnano system coupled to a Q Exactive™ HF Hybrid Quadrupole-Orbitrap™ Mass Spectrometer (ThermoFisher Scientific, UK). Protein digests were loaded onto a trapping column (Acclaim PepMap 100 C18, 75 µm x 2 cm, 3 µm packing material, 100 Å) using 0.1 % (v/v) trifluoroacetic acid, 2 % (v/v) acetonitrile in water at a flow rate of 12 µL min-1 for 7 min. For samples 301, 302 and 304, 5 ng was loaded, and for the Long ALACAT, 303 and 305, 10ng was loaded. The peptides were eluted onto the analytical column (EASY-Spray PepMap RSLC C18, 75 µm x 50 cm, 2 µm packing material, 100 Å) at 30 °C using a linear gradient of 30 minutes rising from 3 % (v/v) acetonitrile/0.1 % (v/v) formic acid (Fisher Scientific, UK) to 40 % (v/v) acetonitrile/0.1 % (v/v) formic acid at a flow rate of 300 nL min^-1^. The column was then washed with 79 % (v/v) acetonitrile/0.1 % (v/v) formic acid for 5 min, and re-equilibrated to starting conditions. The nano-liquid chromatograph was operated under the control of Dionex Chromatography MS Link 2.14.

The nano-electrospray ionisation source was operated in positive polarity under the control of QExactive HF Tune (version 2.5.0.2042), with a spray voltage of 1.8 kV and a capillary temperature of 250 °C. The mass spectrometer was operated in data-dependent acquisition mode. Full MS survey scans between m/z 350-2000 were acquired at a mass resolution of 60,000 (full width at half maximum at m/z 200). For MS, the automatic gain control target was set to 3e^6^, and the maximum injection time was 100 ms. The 16 most intense precursor ions with charge states of 2-5 were selected for MS/MS with an isolation window of 2 m/z units. Product ion spectra were recorded between m/z 200-2000 at a mass resolution of 30,000 (full width at half maximum at m/z 200). For MS/MS, the automatic gain control target was set to 1e^5^, and the maximum injection time was 45 ms. Higher-energy collisional dissociation was performed to fragment the selected precursor ions using a normalised collision energy of 30 %. Dynamic exclusion was set to 30 s.

The raw MS data files were loaded into Thermo Proteome Discoverer v.1.4 (ThermoFisher Scientific, UK) and searched against a custom ALACATs database using Mascot v.2.7 (Matrix Science London, UK) with trypsin as the specified enzyme, one missed cleavage allowed, carbamidomethylation of cysteine, label [^13^C_6_][^15^N_2_]lysine and [^13^C_6_][^15^N_4_]arginine set as fixed modifications and oxidation of methionine set as a variable modification. A precursor mass tolerance of 10 ppm and a fragment ion mass tolerance of 0.01 Da were applied.

