## Supplementary Material for "Synthetic biology meets proteomics: Construction of *à la carte* QconCATs for absolute protein quantification"

### Slide 1
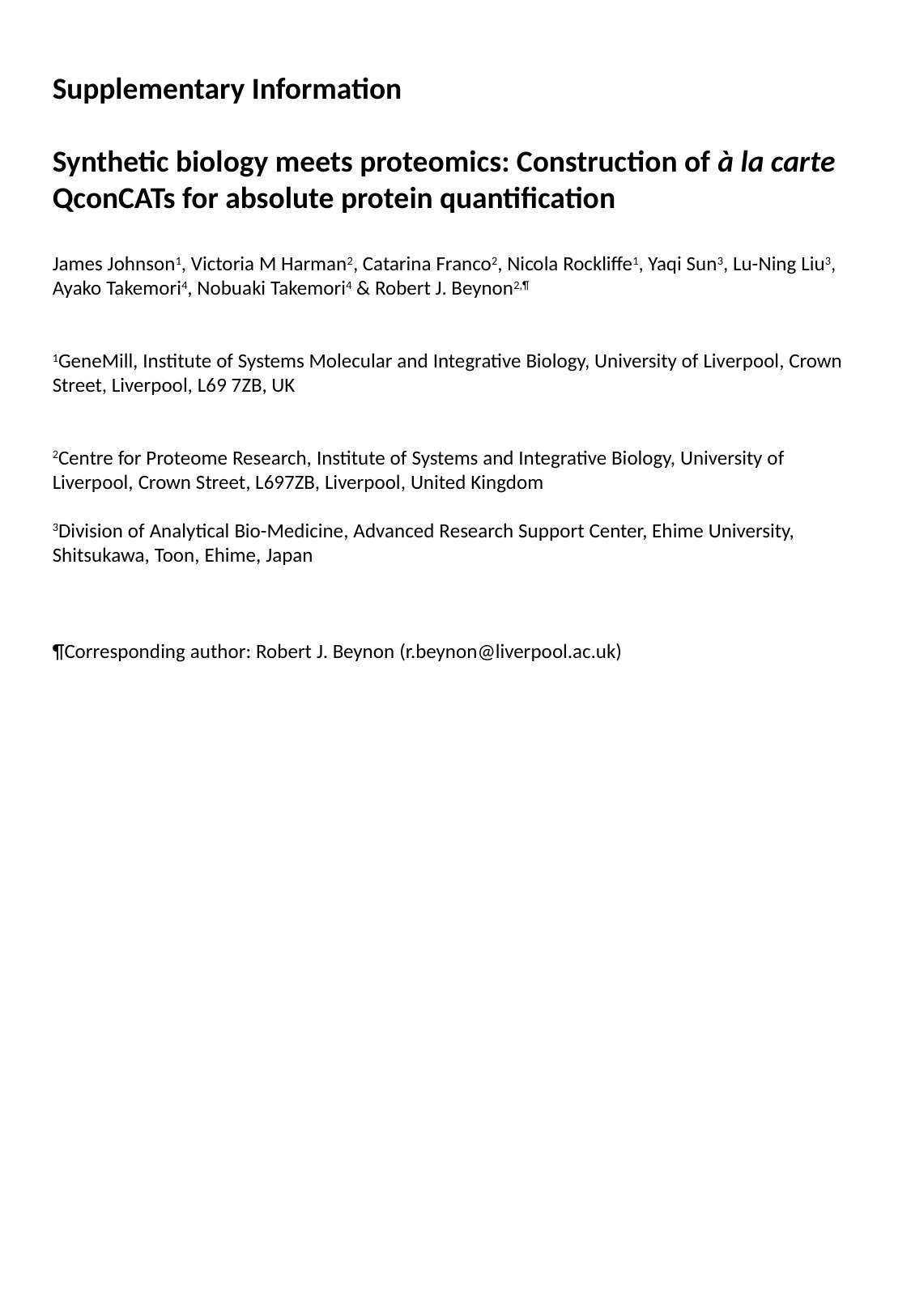

Supplementary Information
Synthetic biology meets proteomics: Construction of à la carte QconCATs for absolute protein quantification
James Johnson^1^, Victoria M Harman^2^, Catarina Franco^2^, Nicola Rockliffe^1^, Yaqi Sun^3^, Lu-Ning Liu^3^, Ayako Takemori^4^, Nobuaki Takemori^4^ & Robert J. Beynon^2,¶^
^1^GeneMill, Institute of Systems Molecular and Integrative Biology, University of Liverpool, Crown Street, Liverpool, L69 7ZB, UK
^2^Centre for Proteome Research, Institute of Systems and Integrative Biology, University of Liverpool, Crown Street, L697ZB, Liverpool, United Kingdom
^3^Division of Analytical Bio-Medicine, Advanced Research Support Center, Ehime University, Shitsukawa, Toon, Ehime, Japan

### Slide 2
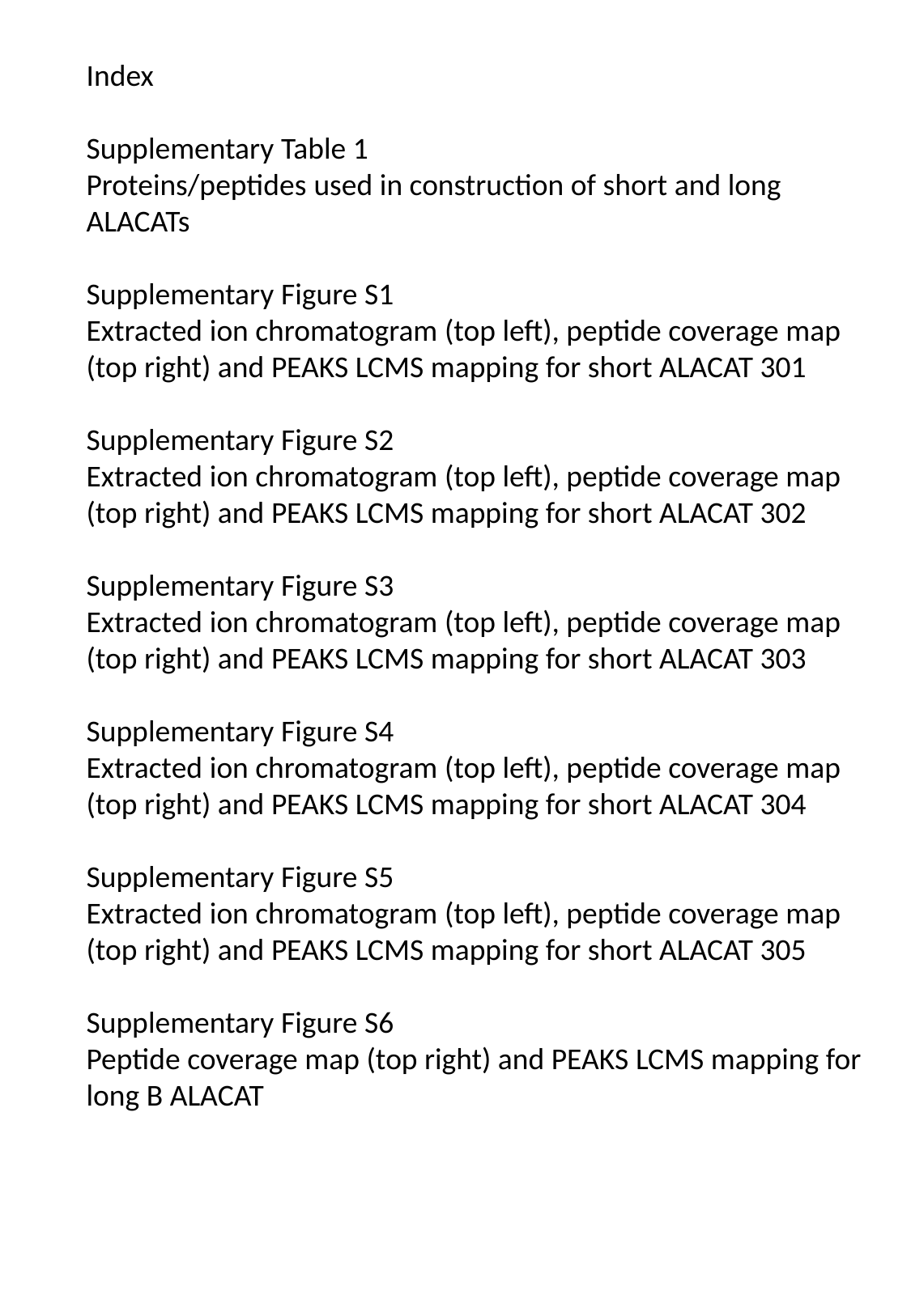

Index
Supplementary Table 1
Proteins/peptides used in construction of short and long ALACATs
Supplementary Figure S1
Extracted ion chromatogram (top left), peptide coverage map (top right) and PEAKS LCMS mapping for short ALACAT 301
Supplementary Figure S2
Extracted ion chromatogram (top left), peptide coverage map (top right) and PEAKS LCMS mapping for short ALACAT 302
Supplementary Figure S3
Extracted ion chromatogram (top left), peptide coverage map (top right) and PEAKS LCMS mapping for short ALACAT 303
Supplementary Figure S4
Extracted ion chromatogram (top left), peptide coverage map (top right) and PEAKS LCMS mapping for short ALACAT 304
Supplementary Figure S5
Extracted ion chromatogram (top left), peptide coverage map (top right) and PEAKS LCMS mapping for short ALACAT 305
Supplementary Figure S6
Peptide coverage map (top right) and PEAKS LCMS mapping for long B ALACAT

### Slide 3
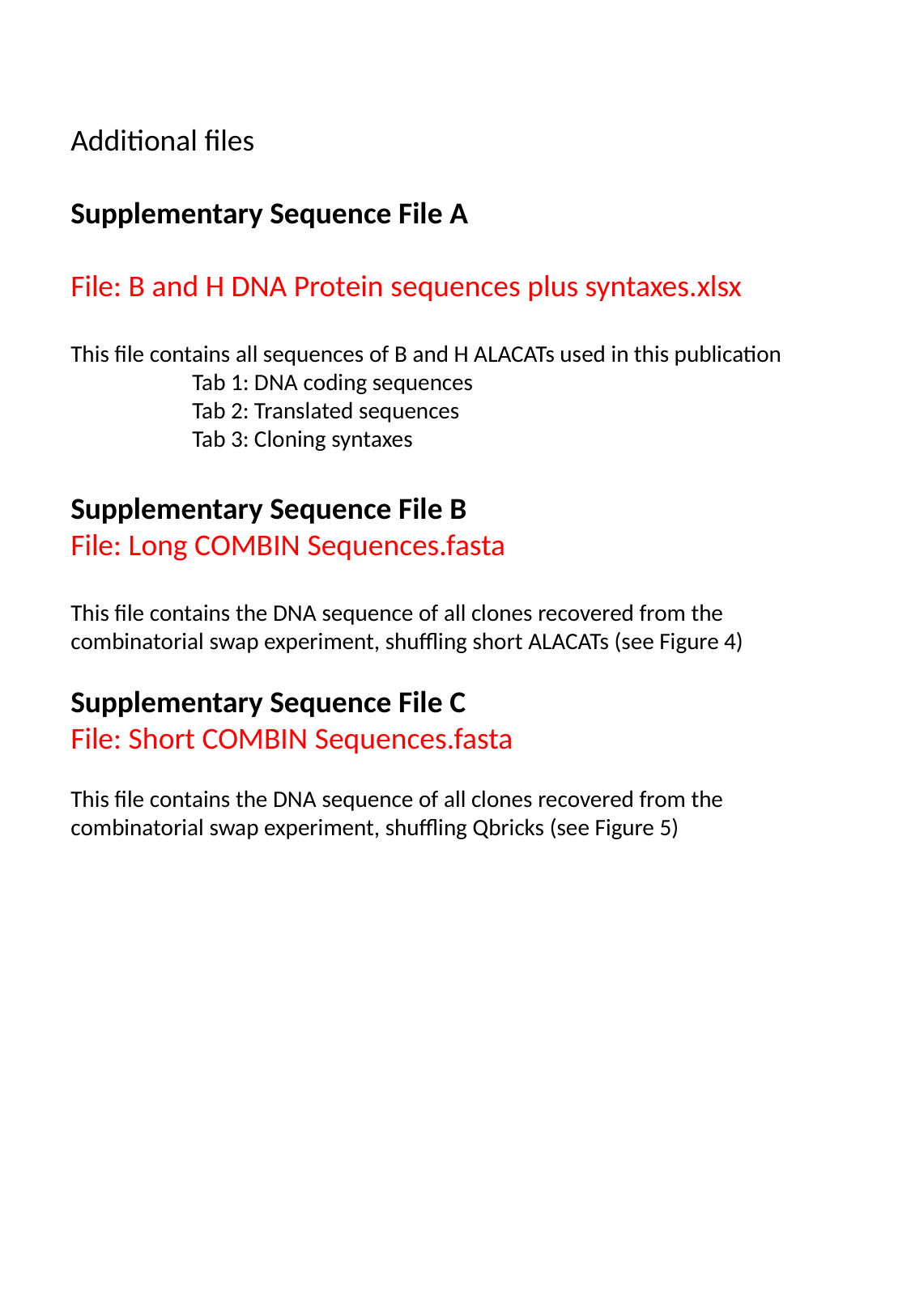

Additional files
Supplementary Sequence File A
File: B and H DNA Protein sequences plus syntaxes.xlsx
This file contains all sequences of B and H ALACATs used in this publication
	Tab 1: DNA coding sequences
	Tab 2: Translated sequences
	Tab 3: Cloning syntaxes
Supplementary Sequence File B
File: Long COMBIN Sequences.fasta
This file contains the DNA sequence of all clones recovered from the combinatorial swap experiment, shuffling short ALACATs (see Figure 4)
Supplementary Sequence File C
File: Short COMBIN Sequences.fasta
This file contains the DNA sequence of all clones recovered from the combinatorial swap experiment, shuffling Qbricks (see Figure 5)

### Slide 4
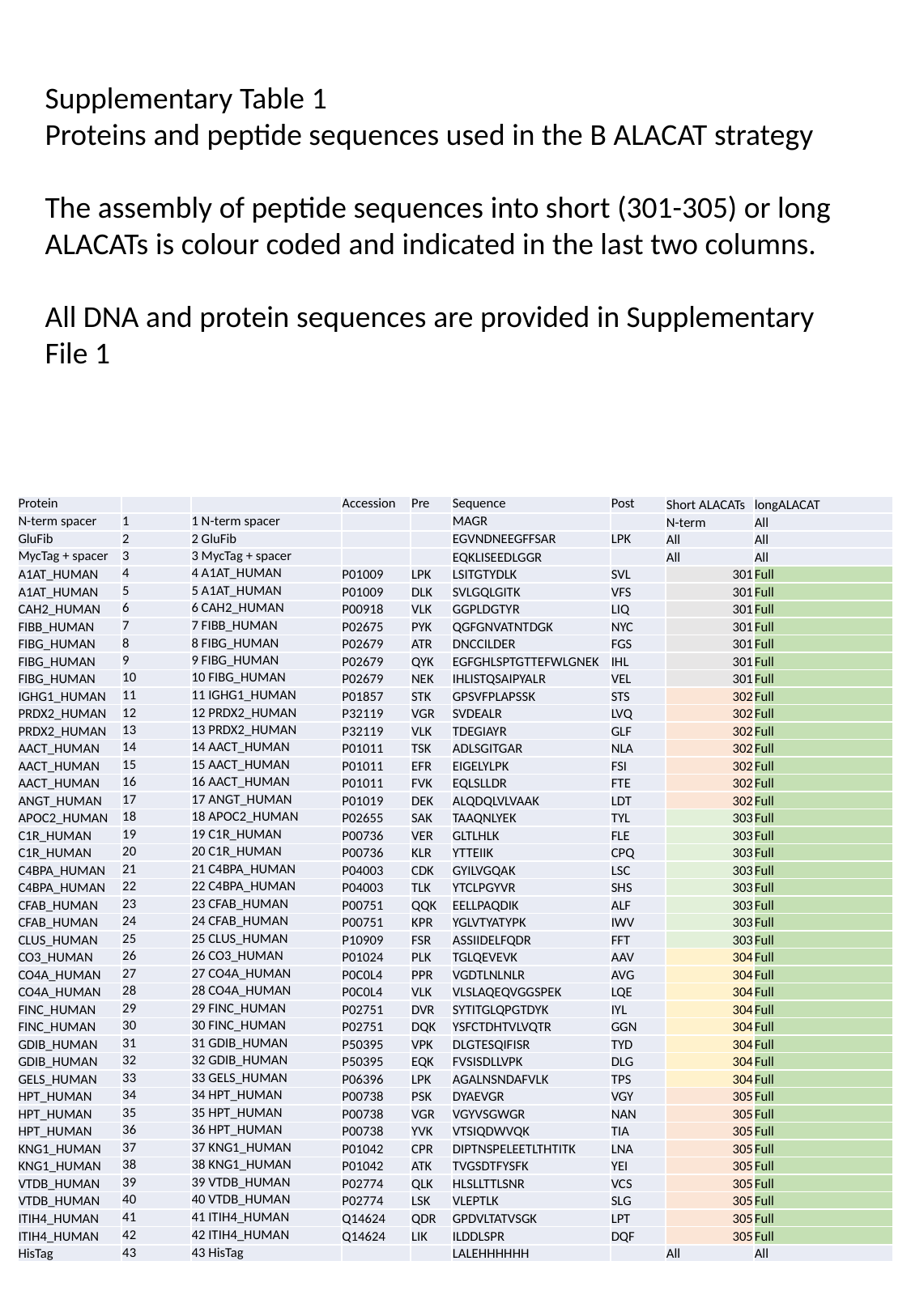

Supplementary Table 1
Proteins and peptide sequences used in the B ALACAT strategy
The assembly of peptide sequences into short (301-305) or long ALACATs is colour coded and indicated in the last two columns.
All DNA and protein sequences are provided in Supplementary File 1
| Protein | | | Accession | Pre | Sequence | Post | Short ALACATs | longALACAT |
| --- | --- | --- | --- | --- | --- | --- | --- | --- |
| N-term spacer | 1 | 1 N-term spacer | | | MAGR | | N-term | All |
| GluFib | 2 | 2 GluFib | | | EGVNDNEEGFFSAR | LPK | All | All |
| MycTag + spacer | 3 | 3 MycTag + spacer | | | EQKLISEEDLGGR | | All | All |
| A1AT\_HUMAN | 4 | 4 A1AT\_HUMAN | P01009 | LPK | LSITGTYDLK | SVL | 301 | Full |
| A1AT\_HUMAN | 5 | 5 A1AT\_HUMAN | P01009 | DLK | SVLGQLGITK | VFS | 301 | Full |
| CAH2\_HUMAN | 6 | 6 CAH2\_HUMAN | P00918 | VLK | GGPLDGTYR | LIQ | 301 | Full |
| FIBB\_HUMAN | 7 | 7 FIBB\_HUMAN | P02675 | PYK | QGFGNVATNTDGK | NYC | 301 | Full |
| FIBG\_HUMAN | 8 | 8 FIBG\_HUMAN | P02679 | ATR | DNCCILDER | FGS | 301 | Full |
| FIBG\_HUMAN | 9 | 9 FIBG\_HUMAN | P02679 | QYK | EGFGHLSPTGTTEFWLGNEK | IHL | 301 | Full |
| FIBG\_HUMAN | 10 | 10 FIBG\_HUMAN | P02679 | NEK | IHLISTQSAIPYALR | VEL | 301 | Full |
| IGHG1\_HUMAN | 11 | 11 IGHG1\_HUMAN | P01857 | STK | GPSVFPLAPSSK | STS | 302 | Full |
| PRDX2\_HUMAN | 12 | 12 PRDX2\_HUMAN | P32119 | VGR | SVDEALR | LVQ | 302 | Full |
| PRDX2\_HUMAN | 13 | 13 PRDX2\_HUMAN | P32119 | VLK | TDEGIAYR | GLF | 302 | Full |
| AACT\_HUMAN | 14 | 14 AACT\_HUMAN | P01011 | TSK | ADLSGITGAR | NLA | 302 | Full |
| AACT\_HUMAN | 15 | 15 AACT\_HUMAN | P01011 | EFR | EIGELYLPK | FSI | 302 | Full |
| AACT\_HUMAN | 16 | 16 AACT\_HUMAN | P01011 | FVK | EQLSLLDR | FTE | 302 | Full |
| ANGT\_HUMAN | 17 | 17 ANGT\_HUMAN | P01019 | DEK | ALQDQLVLVAAK | LDT | 302 | Full |
| APOC2\_HUMAN | 18 | 18 APOC2\_HUMAN | P02655 | SAK | TAAQNLYEK | TYL | 303 | Full |
| C1R\_HUMAN | 19 | 19 C1R\_HUMAN | P00736 | VER | GLTLHLK | FLE | 303 | Full |
| C1R\_HUMAN | 20 | 20 C1R\_HUMAN | P00736 | KLR | YTTEIIK | CPQ | 303 | Full |
| C4BPA\_HUMAN | 21 | 21 C4BPA\_HUMAN | P04003 | CDK | GYILVGQAK | LSC | 303 | Full |
| C4BPA\_HUMAN | 22 | 22 C4BPA\_HUMAN | P04003 | TLK | YTCLPGYVR | SHS | 303 | Full |
| CFAB\_HUMAN | 23 | 23 CFAB\_HUMAN | P00751 | QQK | EELLPAQDIK | ALF | 303 | Full |
| CFAB\_HUMAN | 24 | 24 CFAB\_HUMAN | P00751 | KPR | YGLVTYATYPK | IWV | 303 | Full |
| CLUS\_HUMAN | 25 | 25 CLUS\_HUMAN | P10909 | FSR | ASSIIDELFQDR | FFT | 303 | Full |
| CO3\_HUMAN | 26 | 26 CO3\_HUMAN | P01024 | PLK | TGLQEVEVK | AAV | 304 | Full |
| CO4A\_HUMAN | 27 | 27 CO4A\_HUMAN | P0C0L4 | PPR | VGDTLNLNLR | AVG | 304 | Full |
| CO4A\_HUMAN | 28 | 28 CO4A\_HUMAN | P0C0L4 | VLK | VLSLAQEQVGGSPEK | LQE | 304 | Full |
| FINC\_HUMAN | 29 | 29 FINC\_HUMAN | P02751 | DVR | SYTITGLQPGTDYK | IYL | 304 | Full |
| FINC\_HUMAN | 30 | 30 FINC\_HUMAN | P02751 | DQK | YSFCTDHTVLVQTR | GGN | 304 | Full |
| GDIB\_HUMAN | 31 | 31 GDIB\_HUMAN | P50395 | VPK | DLGTESQIFISR | TYD | 304 | Full |
| GDIB\_HUMAN | 32 | 32 GDIB\_HUMAN | P50395 | EQK | FVSISDLLVPK | DLG | 304 | Full |
| GELS\_HUMAN | 33 | 33 GELS\_HUMAN | P06396 | LPK | AGALNSNDAFVLK | TPS | 304 | Full |
| HPT\_HUMAN | 34 | 34 HPT\_HUMAN | P00738 | PSK | DYAEVGR | VGY | 305 | Full |
| HPT\_HUMAN | 35 | 35 HPT\_HUMAN | P00738 | VGR | VGYVSGWGR | NAN | 305 | Full |
| HPT\_HUMAN | 36 | 36 HPT\_HUMAN | P00738 | YVK | VTSIQDWVQK | TIA | 305 | Full |
| KNG1\_HUMAN | 37 | 37 KNG1\_HUMAN | P01042 | CPR | DIPTNSPELEETLTHTITK | LNA | 305 | Full |
| KNG1\_HUMAN | 38 | 38 KNG1\_HUMAN | P01042 | ATK | TVGSDTFYSFK | YEI | 305 | Full |
| VTDB\_HUMAN | 39 | 39 VTDB\_HUMAN | P02774 | QLK | HLSLLTTLSNR | VCS | 305 | Full |
| VTDB\_HUMAN | 40 | 40 VTDB\_HUMAN | P02774 | LSK | VLEPTLK | SLG | 305 | Full |
| ITIH4\_HUMAN | 41 | 41 ITIH4\_HUMAN | Q14624 | QDR | GPDVLTATVSGK | LPT | 305 | Full |
| ITIH4\_HUMAN | 42 | 42 ITIH4\_HUMAN | Q14624 | LIK | ILDDLSPR | DQF | 305 | Full |
| HisTag | 43 | 43 HisTag | | | LALEHHHHHH | | All | All |

### Slide 5
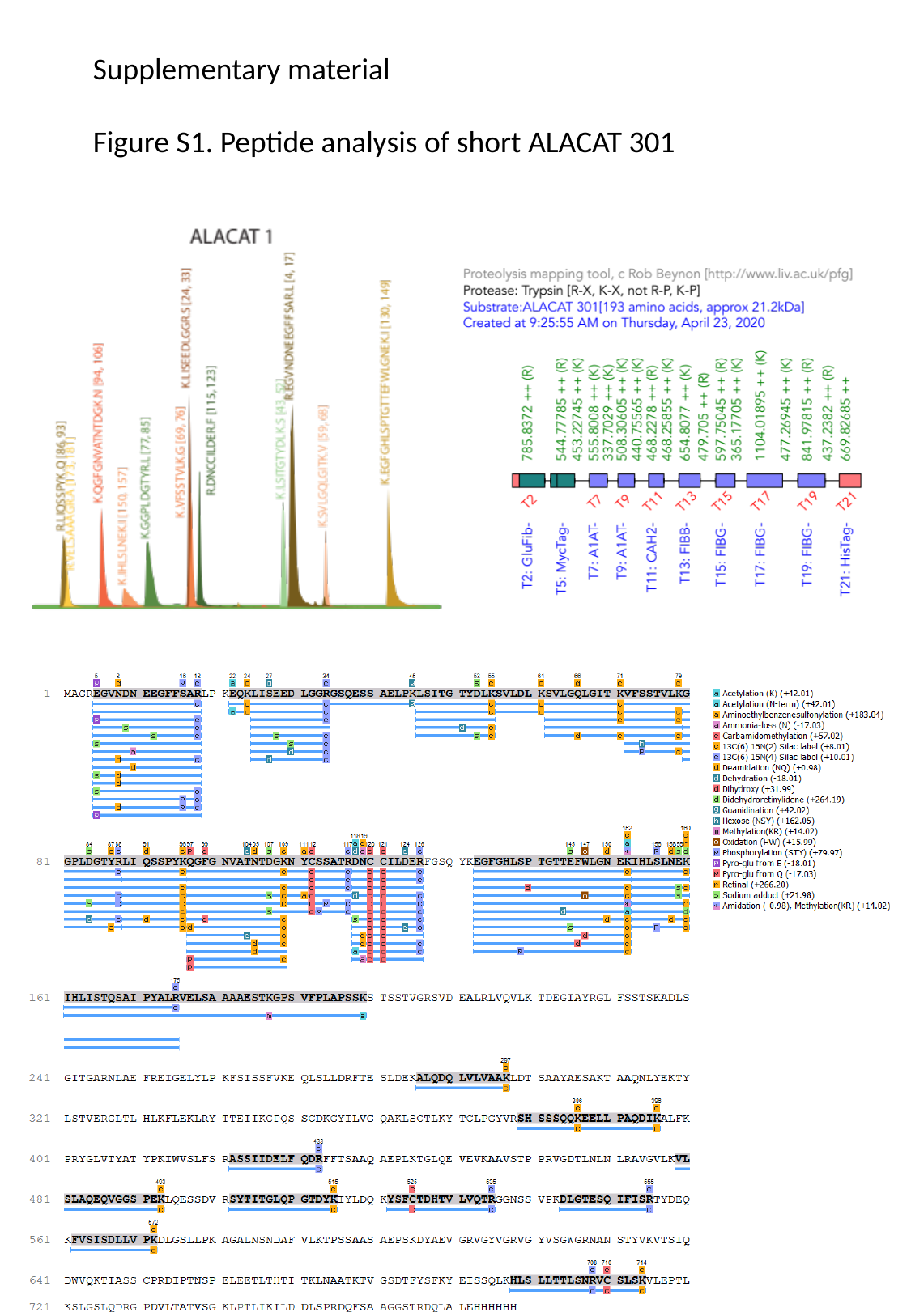

Supplementary material
Figure S1. Peptide analysis of short ALACAT 301

### Slide 6
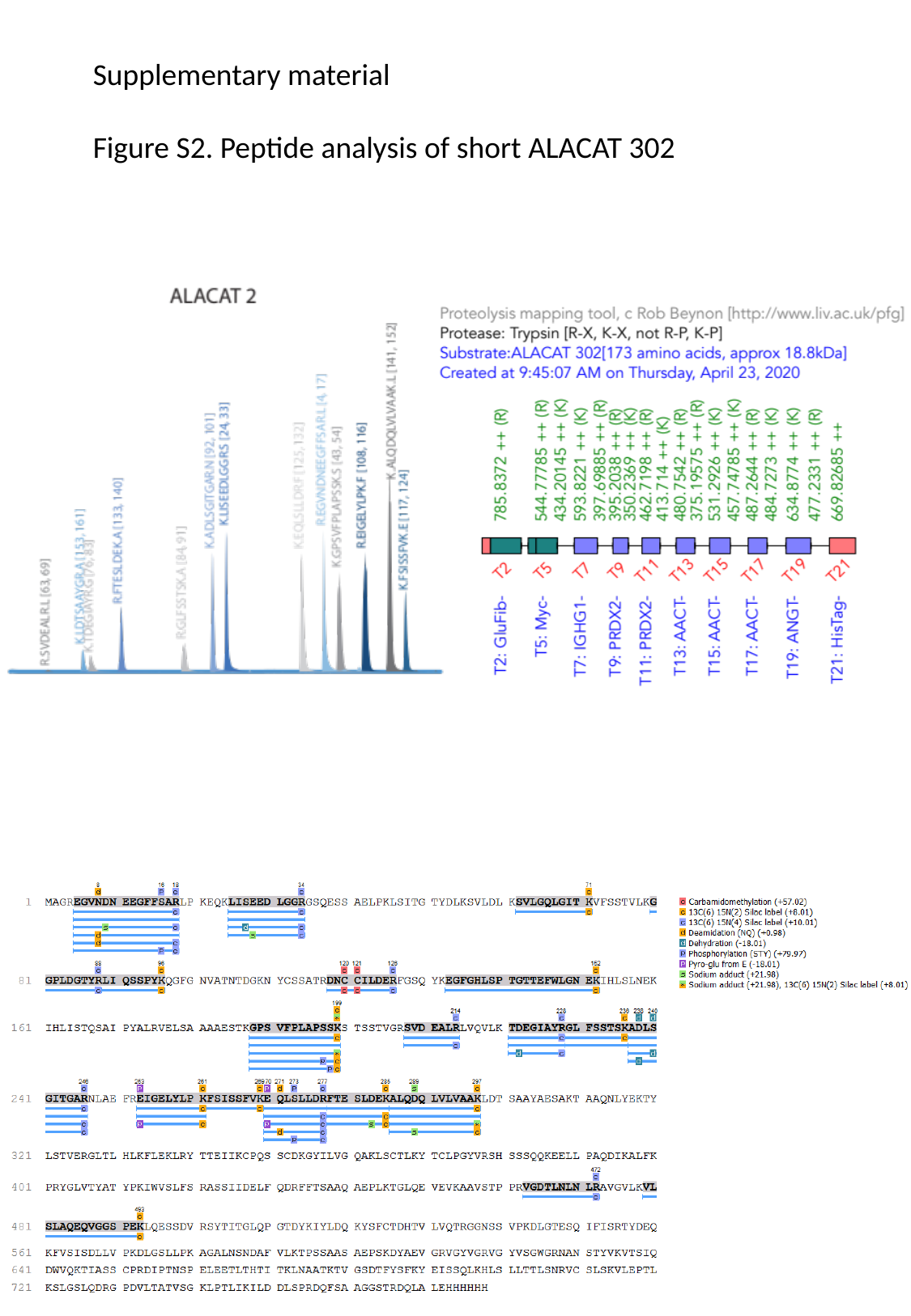

Supplementary material
Figure S2. Peptide analysis of short ALACAT 302

### Slide 7
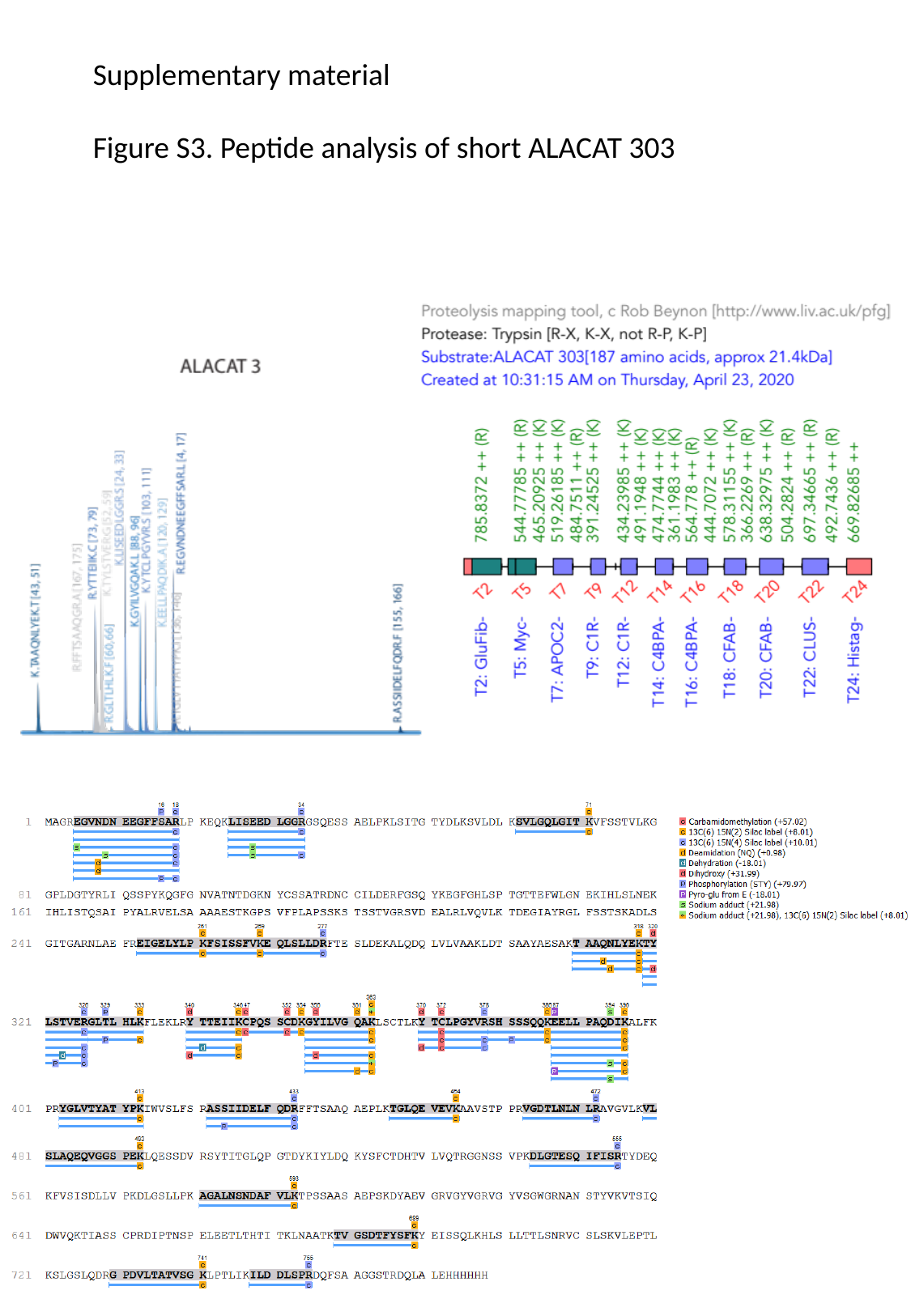

Supplementary material
Figure S3. Peptide analysis of short ALACAT 303

### Slide 8
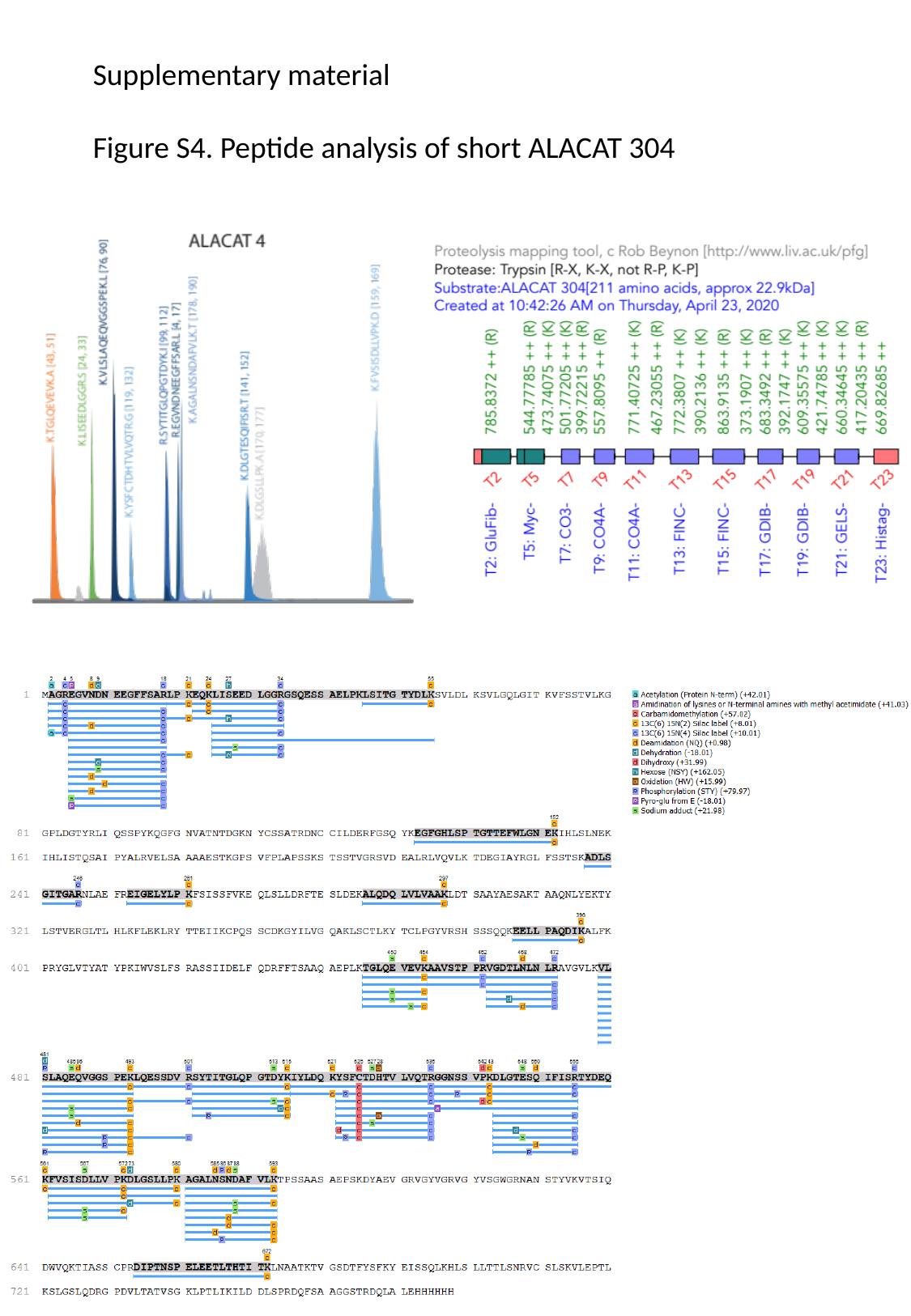

Supplementary material
Figure S4. Peptide analysis of short ALACAT 304

### Slide 9
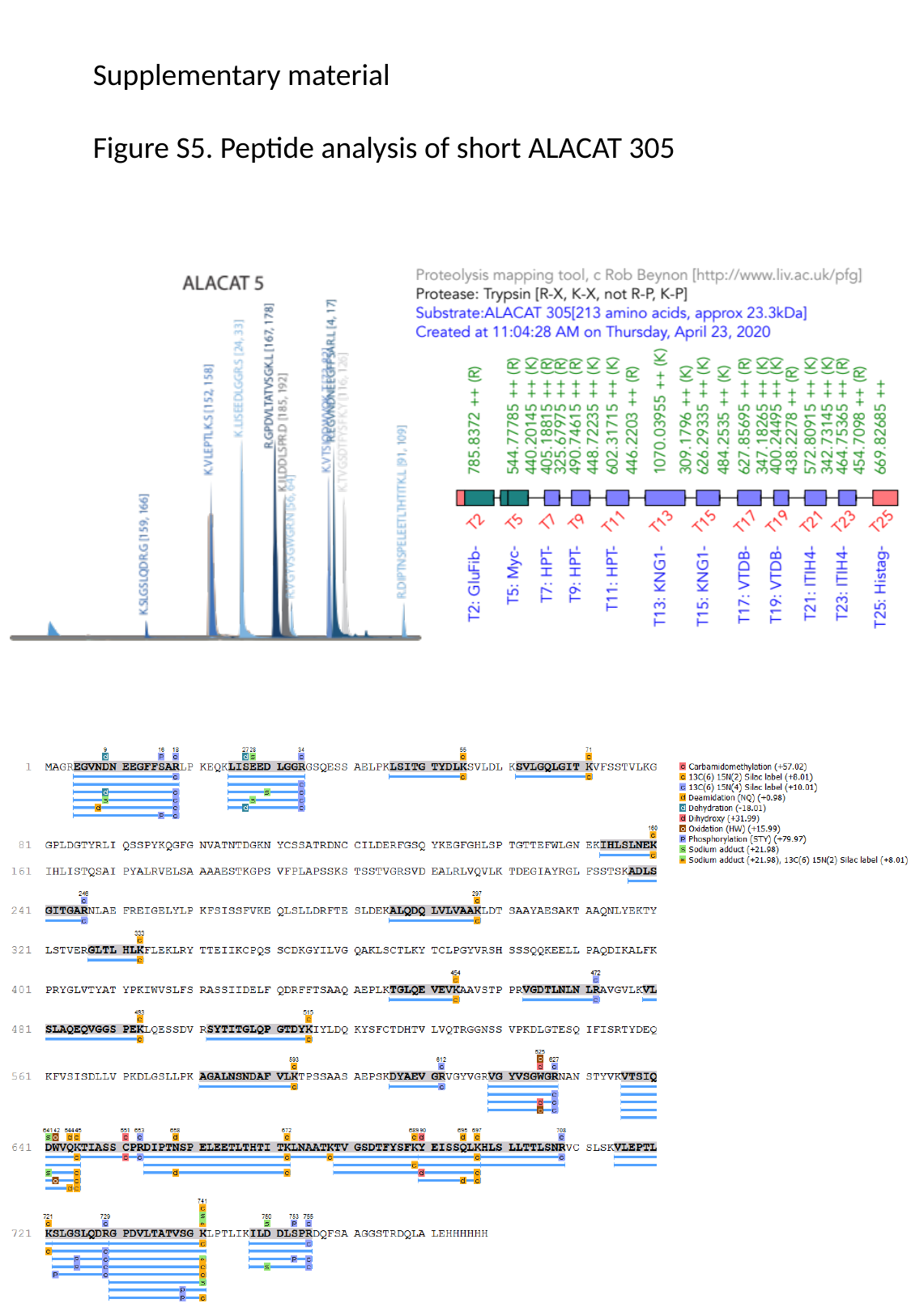

Supplementary material
Figure S5. Peptide analysis of short ALACAT 305

### Slide 10
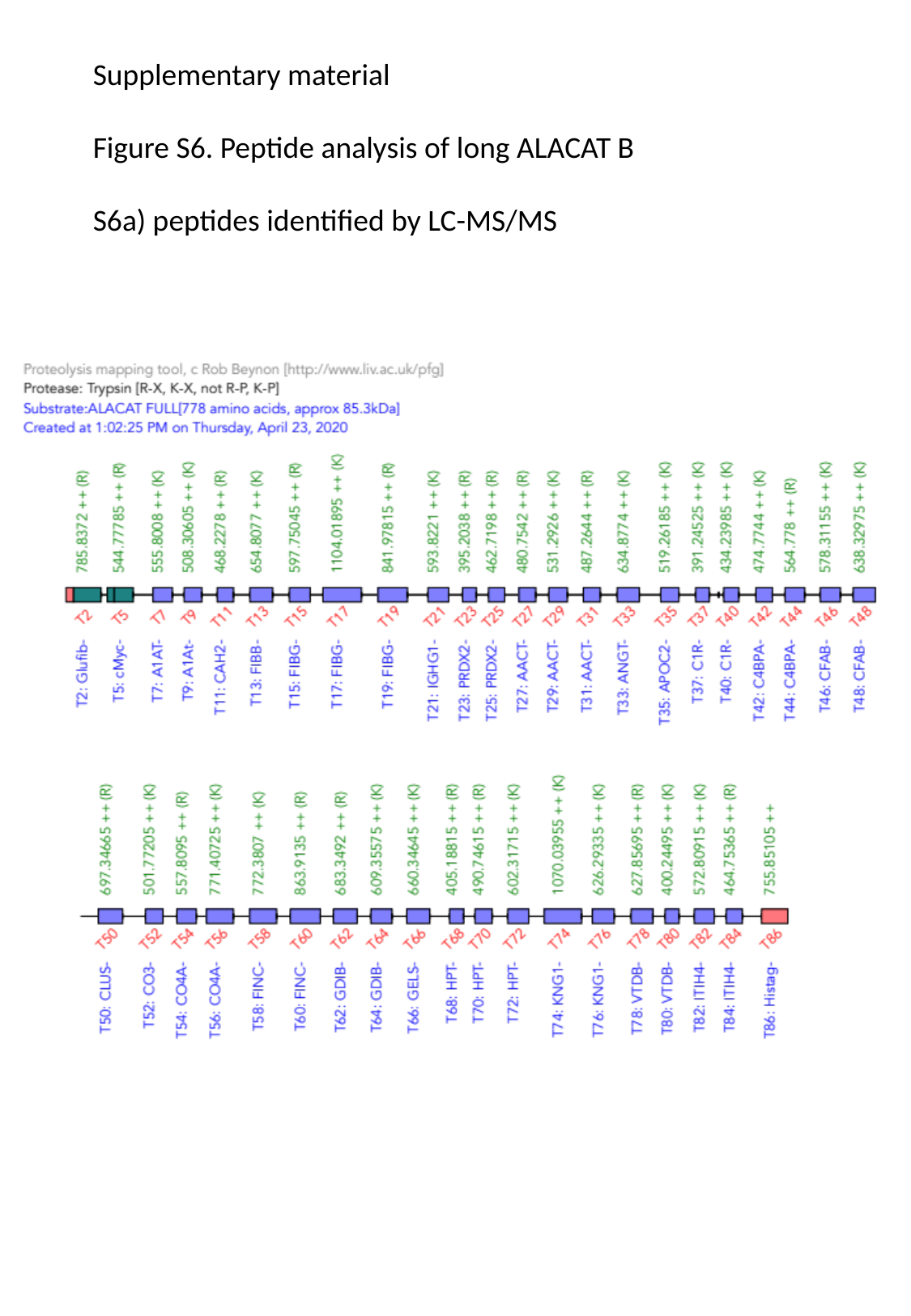

Supplementary material
Figure S6. Peptide analysis of long ALACAT B
S6a) peptides identified by LC-MS/MS

### Slide 11
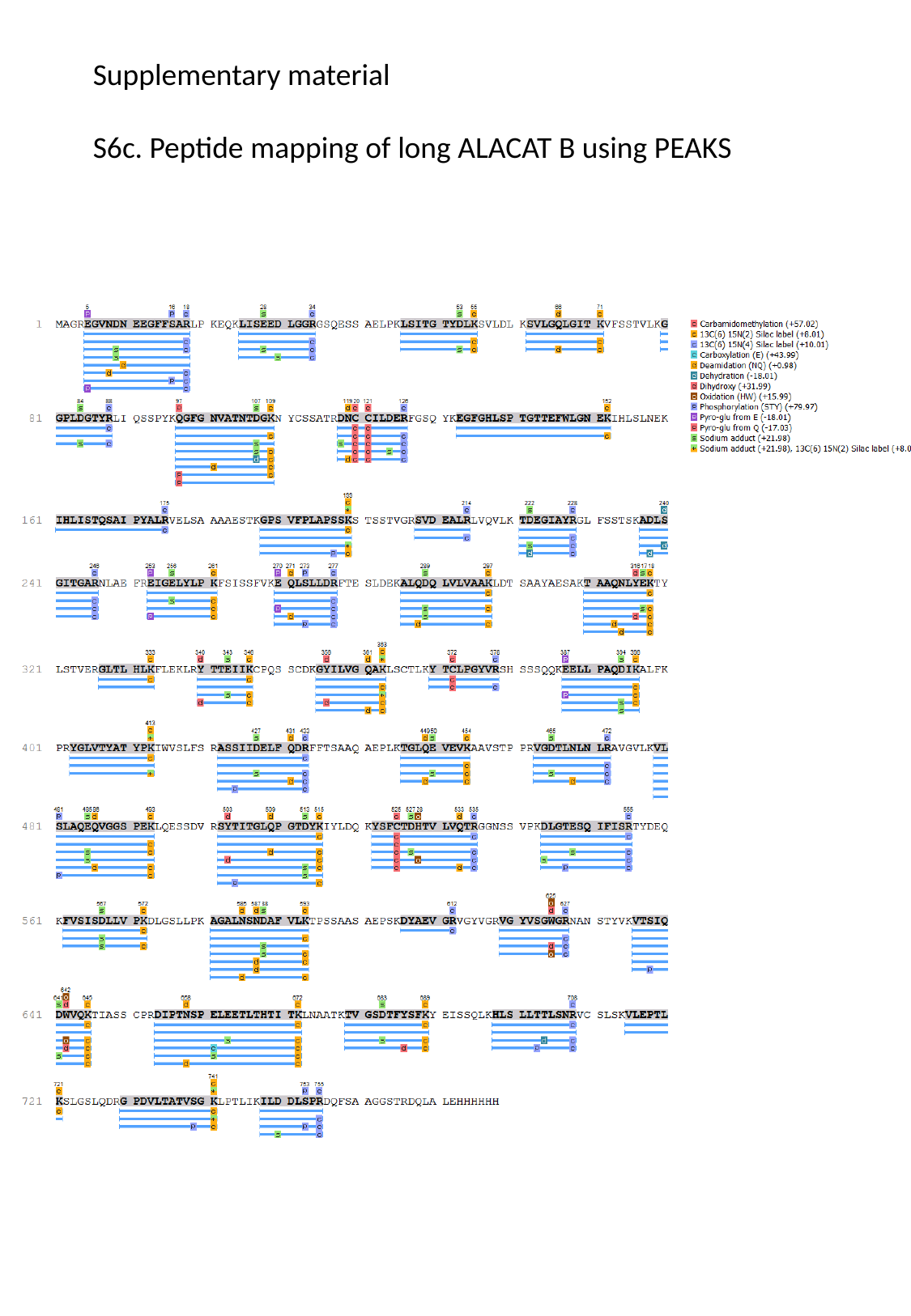

Supplementary material
S6c. Peptide mapping of long ALACAT B using PEAKS
